# Co-expression networks from gene expression variability between genetically identical seedlings can reveal novel regulatory relationships

**DOI:** 10.1101/2020.06.15.152314

**Authors:** S Cortijo, M Bhattarai, J C W Locke, S E Ahnert

## Abstract

Co-expression networks are a powerful tool to understand gene regulation. They have been used to identify new regulation and function of genes involved in plant development and their response to the environment. Up to now, co-expression networks have been inferred using transcriptomes generated on plants experiencing genetic or environmental perturbation, or from expression time series. We propose a new approach by showing that co-expression networks can be constructed in the absence of genetic and environmental perturbation, for plants at the same developmental stage. For this we used transcriptomes that were generated from genetically identical individual plants that were grown in the same conditions and for the same amount of time. Twelve time points were used to cover the 24h light/dark cycle. We used variability in gene expression between individual plants of the same time point to infer a co-expression network. We show that this network is biologically relevant and use it to suggest new gene functions and to identify new targets for the transcription factors GI, PIF4 and PRR5. Moreover, we find different co-regulation in this network based on changes in expression between individual plants, compared to the usual approach requiring environmental perturbation. Our work shows that gene co-expression networks can be identified using variability in gene expression between individual plants, without the need for genetic or environmental perturbations. It will allow further exploration of gene regulation in contexts with subtle differences between plants, which could be closer to what individual plants in a population might face in the wild.

**Author summary:** Plant development and response to changes in the environment are strongly regulated at the level of gene expression. That is why understanding how gene expression is regulated is key, and transcriptome approaches have allowed the analysis of transcription for all genes of the genome. Extracting useful information from the high amount of data generated by transcriptomes is a challenge, and gene co-expression networks are a powerful tool to do this. The principle is to find genes that co-vary in expression in different conditions and to pair them together. Communities of genes that are more closely linked are then identified and this is the starting point to look for their implication in the same pathway. Co-expression networks have been used to identify new regulation and function of genes involved in plant development and their response to the environment. They were constructed using transcriptomes generated on plants experiencing genetic or environmental perturbation. We show that co-expression networks can in fact be constructed in the absence of genetic and environmental perturbation. Our work will allow further exploration of gene co-regulation in contexts with subtle differences between plants, which could be closer to what individual plants in a population might face in the wild.

## Introduction

Understanding how transcriptomes are regulated is key to shedding light on how plants develop and also respond to environmental fluctuations. A powerful tool often used to reveal transcriptional regulation at a genome wide level is gene co-expression networks [1,2]. In gene co-expression networks, genes that co-vary in expression in different conditions are detected and paired together [3–5]. By doing this for the entire transcriptome, a multitude of genes can be linked, indicating a similar gene regulation. Communities of genes, called modules, that are more closely linked can then be identified [6]. The presence of genes in a given module indicates a close co-regulation and is usually the starting point to look for their implication in the same pathway, or their regulation by the same transcription factor(s) [7–10]. Most studies using co-expression networks can be separated into two categories: targeted analyses that use only a subset of genes (selected based on their function or transcriptional regulation) or specific genetic/environmental perturbations, and global analyses that make use of hundreds or thousands of transcriptomes performed in various conditions, often publicly available ones, and do not select genes based on their function prior to the co-expression analysis. Co-expression networks are now commonly used in a variety of work in plant research, and have allowed the identification or prediction of new genes and transcription factors involved in development [9,11,12], in metabolic pathways [13] and in response to biotic and abiotic stresses [8,14–17]. It has also been proposed that the topology of the co-expression network and position of genes in the network can be of interest in itself to identify genes involved in natural diversity in development and in the response to environment [18,19].

One limit of gene co-expression networks is that they only provide information about correlation in expression but do not indicate the direction and type of relationship between genes that are co-expressed. In order to define which genes are transcription factors (TF) that regulate the expression of other genes in the network, additional types of data should be used or integrated [20]. This additional data can be for example ChIP-seq [21,22] that provides the list of targets of a given transcription factor, protein-protein interaction [21,23], as well as the presence of TF binding motifs in the promoter of genes [24,25]. Another limit is that genes should exhibit changes in expression between the different samples used for the analysis in order to detect co-expressed pairs of genes. This is usually achieved by using genetic and/or environmental perturbations in order to cause changes in the transcriptome regulation. However these perturbations often have large effects, and it can be time consuming and challenging to produce the large number of samples required. In order to analyse gene regulation in a biologically context that is more relevant, more subtle changes in expression might be prefered. This could be achieved by using milder genetic or environmental perturbations. Another option would be to analyse changes in expression that occur in the absence of any genetic or environmental perturbation [26]. This can be possible in theory as widespread differences in gene expression levels have been observed between genetically identical plants, in the absence of any environmental perturbation [27–31]. The idea is to use this variability in gene expression to find potential co-regulation. In mammals, variability in gene expression between single cells of the same cell type has been used to identify co-expression patterns for genes that show a high level of gene expression variability between cells [32]. Moreover, gene co-expression networks have been inferred using transcriptomes of individual plant leaves, after removing in silico the genotype, environment and genotype x environment effects on gene expression [26]. The modules identified in this network were functionally relevant and this study allowed the identification of a new regulator of the jasmonate pathway [26]. It thus shows that the analysis of gene expression regulation can be as powerful in the absence of genetic and environmental fluctuation. However, the first step of the study of Bhosale and colleagues was to remove *in silico* the genotype, environment and genotype x environment effects on gene expression, as the transcriptomes were performed on plants from different genotypes, as well as plants that were grown in different research laboratories. It is thus not clear if co-expression networks can be identified in plants from transcriptomes performed in the absence of genetic and environmental perturbation.

We thus decided to test if it is possible to infer gene co-expression networks using transcriptomes generated on single plants in the absence of any genetic and environmental perturbation. In particular, we wanted to define if such a network would provide different information compared to a network using environmental perturbation. Finally we wished to determine if modules that would be detected in such a network would have functional relevance. In order to answer these questions, we took advantage of the existence of a set of published transcriptomes carried out on single seedlings of the same genotype that were grown in the same environmental conditions [28]. In this dataset, multiple genetically identical seedlings had been harvested at several time points during a day/night cycle. Differences in expression between seedlings were previously observed for many genes in each time point of this dataset. In particular, 8.7% of the genes in this dataset have been identified as Highly Variable Genes (HVG), as their expression was statistically more variable between seedlings than the rest of the transcriptome. Using this dataset we were able to infer co-expression networks in absence of genetic and environmental perturbations. Based on enrichment in a module for genes involved in flavonoid metabolism, we speculated that AT4G22870, a 2-oxoglutarate (2OG) and Fe(II)-dependent oxygenase, could also have a role in flavonoid metabolism. Finally, we identified new targets for the TFs PHYTOCHROME INTERACTING FACTOR 4 (PIF4), GIGANTEA (GI) and PSEUDO-RESPONSE REGULATOR 5 (PRR5).

## Results

### Co-expression networks can be inferred using expression variation between individual seedlings

Co-expression networks in plants are normally inferred using transcriptomes obtained from pools of plants, using genetic or environmental perturbations in order to identify genes that co-vary in expression between these conditions. In order to define if co-expression networks can be inferred from expression measurements obtained from single seedlings in the absence of genetic and environmental perturbations, we used the previously published dataset of transcriptomes generated on single seedlings grown in the same environment. This dataset contained a total of fourteen seedlings per time point, for twelve time points spanning a 24 hours day/night cycle [28]. Widespread differences in expression levels have been observed between seedlings in this dataset, which is a prerequisite to be able to infer a co-expression network ([3–5] Fig 1a and S1). We first detected co-expressed genes in each time point, by measuring Spearman correlation for each pair of genes in profiles of expression in the 14 seedlings of this time point. In order to keep robust correlations in the final network, we then selected edges of the network that are detected in at least four consecutive time points, with one gap allowed (Materials and Methods). Using this approach, we find a total of 4715 edges, connecting 1729 genes in this network, from now on referred to as the variability network. The number of edges detected for each time point varies from 787 to 3221, with a higher number of edges being detected at the end of the day and the beginning of the night (Fig 1b). We then used the Louvain community detection algorithm in order to identify modules of genes that are densely connected in the network [33]. In total, we identified 153 modules (Table S1), containing between 2 and 334 genes, with most of the modules only composed of two genes (Fig 1c). To test the robustness of the variability network, we also selected edges that are in at least 3 consecutive time points and compared the detected modules in both networks (Fig S2). Similar modules with a similar overall connectivity between them are found in the two networks, which confirms the robustness of the modules in our original network. Modules in the network based on 4 consecutive time points are smaller. In somes cases, several modules of the network based on 4 consecutive time points correspond to a single module in the network based on 3 consecutive time points and these smaller modules have differences in several features (Fig S2). That is why we decided to focus our analysis on the network obtained when selecting edges present in 4 consecutive time points, and in particular for the 28 modules of this network containing 5 genes or more.

**Fig 1.**
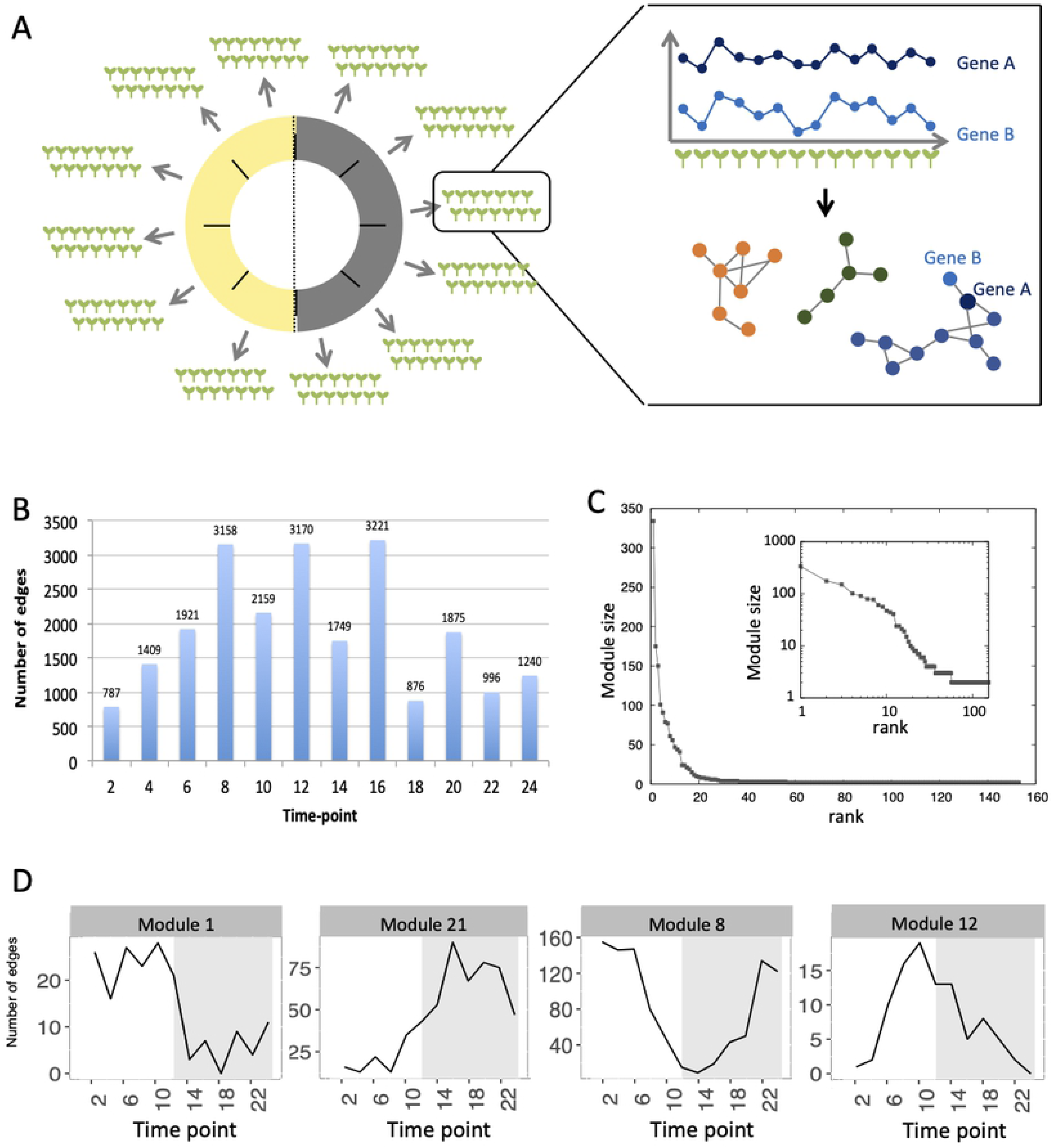
Inference of gene co-expression networks in absence of genetic and environmental perturbations. (A) Description of co-expression network inference using transcriptomes performed on single seedlings. Transcriptomes were generated for a total of fourteen seedlings per time point, with twelve time points spanning a day/night cycle over 24 hours. In each time point, genes with correlated expression profiles in the fourteen seedlings were identified. The co-expression network was inferred based on pairs of genes correlated for at least four consecutive time points. Finally, modules in the network, which consist of groups of genes that are densely connected, were detected. (B) Total number of edges in the final network that are detected in each time point. (C) Distribution of the number of genes present in each of the 153 modules. Inset shows the same data plotted with a logarithmic scale. (D) Number of edges that are detected in each time point for four modules: the module 1 in which most edges are detected during day time, the module 21 in which most edges are detected during the night time, the module 8 in which most edges are detected at the transition between night and day and the module 12 in which most edges are detected at the transition between day and night.

First we analysed the number of edges at each time point throughout the time course for each module (Fig 1d and S3). In most modules the edges are distributed non-uniformly across the 12 time points. Some exhibit a larger number of edges during the day or the night, while in other modules, a larger number of edges is observed at the transitions from night to day, or from day to night. It indicates that genes in these modules are co-regulated at some moments of the day/night cycle but not at others. While for some modules, this is linked with the genes being more expressed at these same times of the day (module 1 for example), this is not the case for other modules in which genes are expressed throughout the time course (module 12 for example, Fig 2). Most of the edges in module 1 are observed during the day (Fig 1d) and we were able to confirm co-expression of genes in this module by doing a RT-qPCR in a replicate experiment for a few genes in this module. In this replicate experiment, we find a very high correlation during the day (ZT6), and a lower correlation during the night (ZT14) (Fig S4). On the other hand, most edges in module 21 are observed during the night (Fig 1d). We also find in a replicate RT-qPCR experiment that genes of module 21 were more correlated during the night than the day (Fig S4). These results confirm that the co-expression of genes in modules of the variability network, and also the differences in co-expression between the day and night, can be reproduced in a replicate experiment. Moreover, we find that modules with a high percentage of edges during the night are more connected to one another than with modules for which most edges are observed during the day, and vice versa (Fig 5a). We can measure this assortativity of the network (i.e. the tendency of similar nodes to be connected to each other) through the Pearson correlation of the daytime-specific edge percentages of connected modules (Pearson correlation=0.4573, p-value=0.043). This result shows that modules that are more connected to one another are more similar, at least for this feature, indicating that the community detection in the network worked well and provides modules that are relevant.

**Fig 2.**
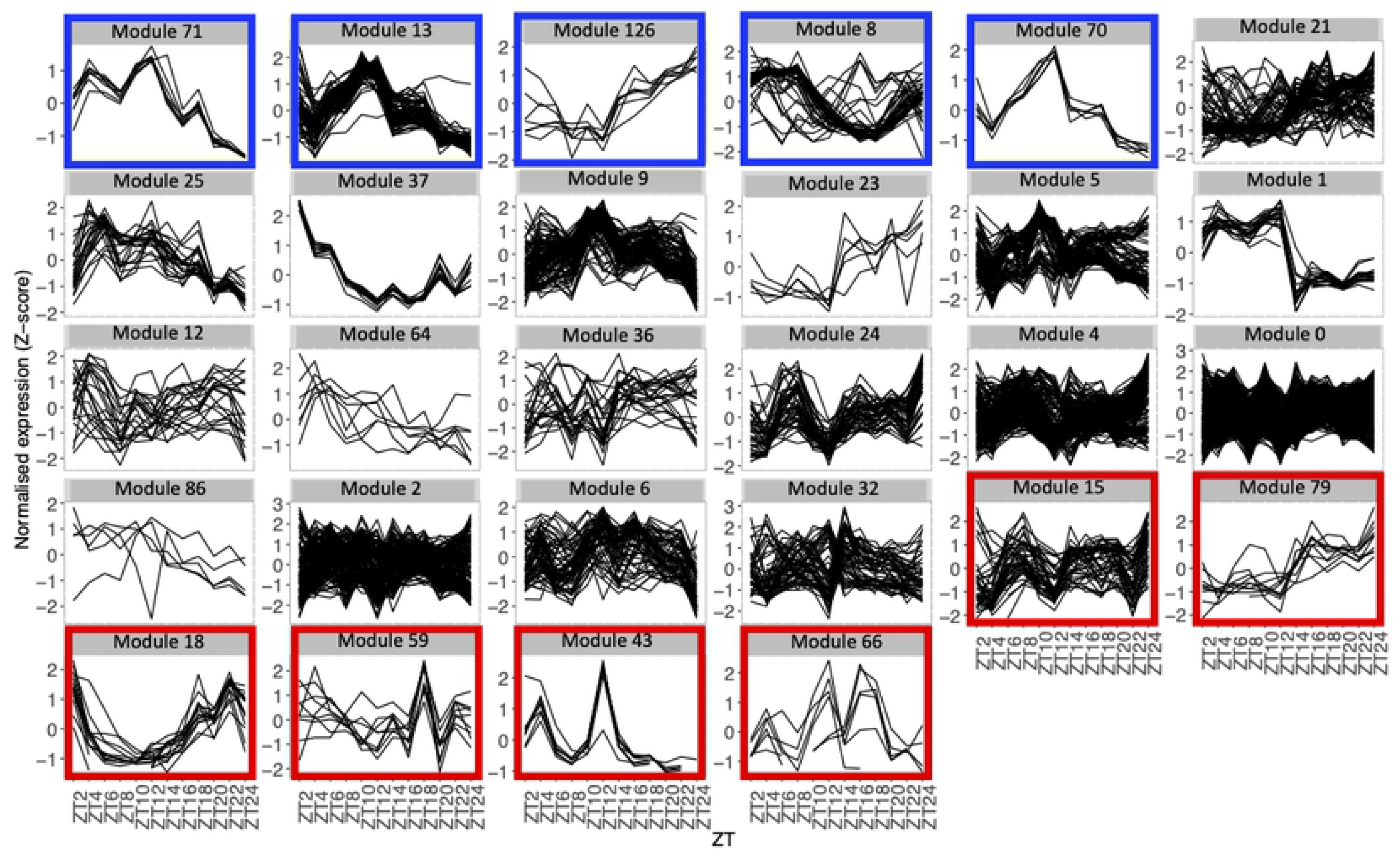
Comparison of the variability network and the averaged time course network. Expression profiles throughout the time course for genes in each module with 5 genes or more, using the average expression of the fourteen seedlings for each time point. Each line represents the normalised expression (z-score) for one gene. Modules are ordered by the percentage of genes in the averaged time course network (high to low). Modules highlighted in blue contain 50% or more of genes that are also in the averaged time course network. Modules highlighted in red contain 15% or less of genes that are also in the averaged time course network.

Since high gene expression variability between genetically identical plants was previously observed in the transcriptome dataset we used to infer the variability network [28], we tested if the network is enriched in highly variable genes (HVGs). We find a total of 477 HVGs in the network, that is 27.6% of all genes in the variability network. This is higher than the 8.7% of HVGs that were detected in the full transcriptome dataset [28]. This result suggests that most of the genes in the variability network do not have to display a high level of gene expression variability to be able to detect co-expression between individual seedlings. We find that most modules are either strongly enriched in HVGs, or strongly depleted in HVGs, with only a few modules containing around 27% of HVGs (Table 1, Fig S5a). Modules 37, 43 and 66 for example are only composed of HVGs, while a total of 8 modules do not have a single HVG. This result suggests that HVGs can co-vary in expression and are potentially co-regulated. It also suggests that HVGs are not likely to co-vary in expression with non variable genes. To test if this result could indicate a bias in the method used to construct the variability network and detect modules, we analysed expression levels in single seedlings for genes in modules with high or low percentage of HVG (Fig S5b). We find that modules with high or low percentage of HVG have different expression profiles in the seedlings, indicating an absence of bias. Moreover, modules with a high percentage of HVG tend to be more connected to one another than with modules containing a low percentage of HVG, and vice versa (Pearson correlation=0.6896, p-value= 0.0007683, Fig 5b).

**Table 1.** Number and percentage of HVGs in each module, for modules with at least 5 genes.

| module name | number of genes in module | number of HVGs in module | percentage of HVGs in module | Fisher's p-value |
| --- | --- | --- | --- | --- |
| 37 | 12 | 12 | 100,00 | 0,0022 |
| 43 | 8 | 8 | 100,00 | 0,0118 |
| 66 | 6 | 6 | 100,00 | 0,0287 |
| 18 | 15 | 14 | 93,33 | 0,0022 |
| 24 | 47 | 42 | 89,36 | 0,0001 |
| 59 | 9 | 8 | 88,89 | 0,0181 |
| 15 | 44 | 38 | 86,36 | 0,0001 |
| 1 | 19 | 15 | 78,95 | 0,0052 |
| 32 | 56 | 43 | 76,79 | 0,0001 |
| 64 | 7 | 5 | 71,43 | 0,1496 |
| 23 | 6 | 4 | 66,67 | 0,2384 |
| 21 | 79 | 50 | 63,29 | 0,0001 |
| 79 | 10 | 6 | 60,00 | 0,1322 |
| 4 | 175 | 97 | 55,43 | 0,0001 |
| 25 | 24 | 9 | 37,50 | 0,4014 |
| 8 | 41 | 8 | 19,51 | 0,4820 |
| 2 | 150 | 17 | 11,33 | 0,0002 |
| 13 | 77 | 6 | 7,79 | 0,0009 |
| 9 | 101 | 3 | 2,97 | 0,0001 |
| 0 | 334 | 3 | 0,90 | 0,0001 |
| 5 | 91 | 0 | 0,00 | 0,0001 |
| 6 | 61 | 0 | 0,00 | 0,0001 |
| 36 | 24 | 0 | 0,00 | 0,0047 |
| 12 | 21 | 0 | 0,00 | 0,0127 |
| 71 | 8 | 0 | 0,00 | 0,2142 |
| 126 | 7 | 0 | 0,00 | 0,3578 |
| 70 | 6 | 0 | 0,00 | 0,3512 |
| 86 | 5 | 0 | 0,00 | 0,5915 |

Our results show that gene co-expression networks can be inferred in the absence of genetic or environmental perturbation. Moreover, genes don’t need to show a high level of gene expression variability between seedlings to be integrated in the network.

### Additional gene co-expression is identified in the variability network compared to a network inferred from a pool of plants

Next, we decided to test whether the co-expression network based on the variability of expression between genetically identical plants grown in the same environment is different from a co-expression network inferred from a pool of plants. Since the transcriptome dataset contains data for several time points throughout a day/night cycle, we decided to infer a co-expression network using the average expression of the fourteen seedlings for each time point and thus exploit changes in expression happening during the time course. This network, referred to as the averaged time course network, allows the identification of co-expression throughout the time course. Using this approach, we find a total of 9332 edges, connecting 3861 genes in the averaged time course network. A total of 524 genes of this averaged time course network are also present in the variability network, that is 30% of the genes in the variability network (Table 2). Only 35 edges are shared between the two networks. This result shows that the majority of the genes and edges present in the variability network are not detected in this dataset using a classical approach with pools of plants.

**Table 2.** Number and percentage in each module of genes also detected in the averaged time course network.

| module name | number of genes in module | number of genes in module also in averaged time course network | percentage of genes in module also in averaged time course network | Fisher's p-value |
| --- | --- | --- | --- | --- |
| 71 | 8 | 7 | 87,5 | 0,0585 |
| 13 | 77 | 59 | 76,6 | 0,0001 |
| 126 | 7 | 4 | 57,1 | 0,2947 |
| 8 | 41 | 21 | 51,2 | 0,0671 |
| 70 | 6 | 3 | 50,0 | 0,4431 |
| 21 | 79 | 37 | 46,8 | 0,0431 |
| 25 | 24 | 10 | 41,7 | 0,4146 |
| 37 | 12 | 5 | 41,7 | 0,566 |
| 9 | 101 | 41 | 40,6 | 0,1273 |
| 23 | 6 | 2 | 33,3 | 1 |
| 5 | 91 | 29 | 31,9 | 0,8248 |
| 1 | 19 | 6 | 31,6 | 1 |
| 12 | 21 | 6 | 28,6 | 1 |
| 64 | 7 | 2 | 28,6 | 1 |
| 36 | 24 | 6 | 25,0 | 0,8289 |
| 24 | 47 | 11 | 23,4 | 0,5296 |
| 4 | 175 | 37 | 21,1 | 0,0592 |
| 0 | 334 | 69 | 20,7 | 0,0063 |
| 86 | 5 | 1 | 20,0 | 1 |
| 2 | 150 | 26 | 17,3 | 0,0087 |
| 6 | 61 | 10 | 16,4 | 0,0845 |
| 32 | 56 | 9 | 16,1 | 0,0985 |
| 15 | 44 | 6 | 13,6 | 0,0627 |
| 79 | 10 | 1 | 10,0 | 0,4744 |
| 18 | 15 | 1 | 6,7 | 0,1401 |
| 59 | 9 | 0 | 0,0 | 0,1286 |
| 43 | 8 | 0 | 0,0 | 0,2106 |
| 66 | 6 | 0 | 0,0 | 0,3465 |

We find that between 0% and 87.5% of genes in modules of the variability network are also in the averaged time course network, with most of the modules having between 20 and 50% of genes also present in the averaged time course network (Table 2). The modules with the highest percentage of genes also in the averaged time course network are modules 71 (87%: 7 out of 8 genes) and 13 (76%: 59 out of 77 genes).

We find that genes in these modules have very similar and clear expression profiles throughout the time course (Fig 2). This is also the case for all modules with at least 50% of genes in the averaged time course network (modules highlighted in blue in Fig 2). This result could suggest that the reason why these modules contain many genes also present in the averaged time course network, is because their genes have very similar expression patterns throughout the day/night cycle. On the other hand, several modules that only have 15% or less of genes present in the averaged time course network are composed of genes without clear expression patterns during the time course. These results show that additional gene co-expression is identified in the variability network compared to the averaged time course network. Most importantly, using gene expression in single seedlings, co-expression between genes can be detected even in absence of expression patterns throughout the day/night cycle.

### Modules identified in the variability network are functionally relevant

In order to define if the modules identified in the variability network are functionally relevant, we performed a gene ontology (GO) enrichment analysis. We find that some of the modules have strongly enriched GO (Table S2).

For example, the module 8 is enriched in multiple GO related to photosynthesis. In particular, 33 genes out of the 41 in this module are members of the photosystem I or II, or of the light harvesting complex (Fig 3a, Table S3). Other genes in this module also have functions related to photosynthesis: CURT1A is required for a proper thylakoid morphology [34], while RBCS1A, RBCS3B and RCA are members of the Rubisco or necessary to Rubisco light activation [35,36]. Most edges of module 8 are observed at the transition between night and day. This module contains 51 % of genes that are also in the averaged time course network, which could be expected as most of the genes have very similar expression patterns throughout the day/night cycle. In particular, all genes of this module present in the photosystem I, II or the light harvesting system have the same expression profile with a peak of expression at dawn and the beginning of the day, while other genes have different expression profiles with a peak of expression at the beginning of the day and another one during the night (Fig S6a). Also, we find that these other genes are at the periphery of the module 8 (Fig S6b), which highlights that these genes are less well correlated with the dense core of highly correlated photosystem genes in the centre of the network. Another module enriched in GO related with photosynthesis is module 37 (Table S2), in which 9 out of the 12 genes are chloroplast genes, some being present in the Photosystems I or II, in the Cytochrome b6/f complex or in the ATP synthase (Fig 3a, Table S3). Genes in module 37 are mainly expressed at the beginning of the day. These results suggest that the expression of genes involved in photosynthesis are co-regulated, not only over time, but also between plants at a given time.

**Fig 3.**
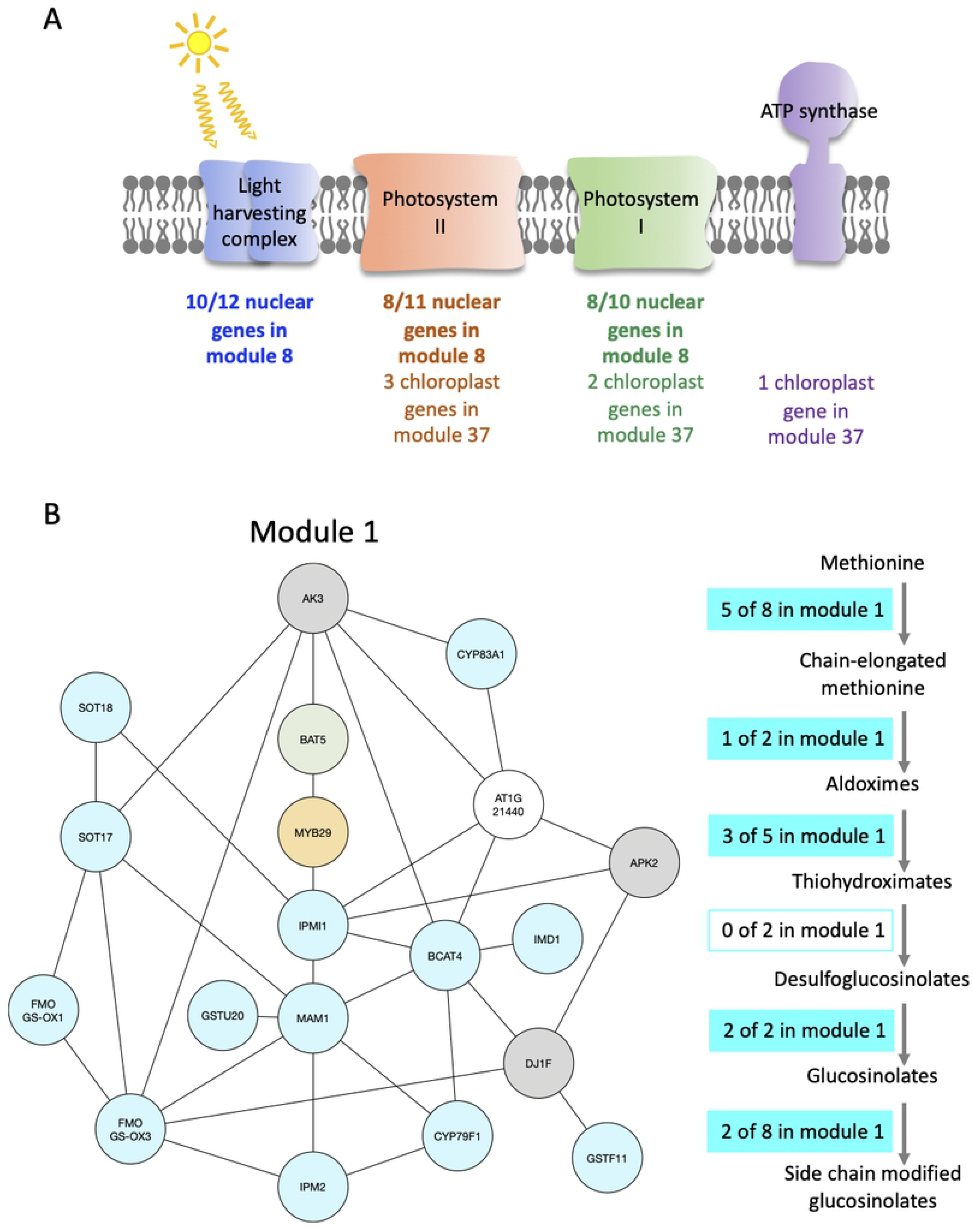
Modules enriched in genes involved in the photosynthesis and the glucosynolate pathway. (A) Functional analysis of modules 8 and 37. For each module, the number of genes that are part of the Photosystem I (green), Photosystem II (orange), the Light harvesting complex (blue) or the ATP synthase (purple) are indicated. (B) Functional analysis of module 1. Genes of the module are color coded depending on their role in the glucosynolate pathway: biosynthesis (turquoise), transport (green) or regulation (orange). Genes previously identified as co-expressed with glucosynolate biosynthesis genes are also indicated (grey). On the right side, the glucosinolate biosynthesis pathway is shown with an indication of the number of genes present in the module 1 at each step of the pathway.

Module 71 is enriched in GO related to DNA packaging (Table S2), and is in fact only composed of histones, including 2 variants of H2A, 2 variants of H2B, and H3.1 (Table S3). None of the genes in this module are HVGs, and 7 genes out of 8 are also present in the averaged time course network.

Module 1 is enriched in GO related to glucosinolate (Table S2). We find that 16 out of 19 genes of the module are in the glucosinolate biosynthesis pathway, transporters of glucosinolate, or transcription factors (TF) regulating the pathway (Fig 3b, Table S3). All genes in module 1, except one, were previously identified as co-expressed, in a previous study of the glucosinolate pathway [13]. Among the genes that are not known to be involved in glucosinolate biosynthesis, but are co-expressed with it, AKN2 is regulated by the MYB TF also regulating the glucosinolate pathway [37]. AKN2 is involved in sulfate assimilation which is linked to glucosinolate metabolism [37]. It thus makes sense that AKN2 is co-expressed with genes of the glucosinolate biosynthesis pathway. Most edges of module 1 are observed during the day, which is when the genes in the module are more expressed. Also 15 out of the 19 genes of the modules are HVGs.

Module 43 is enriched in GO related to flavonoid metabolism (Table S2), with 6 out of 8 genes shown to be involved in flavonoid biosynthesis [38]. Among the other genes, AT4G22870 has not been shown to be involved in the flavonoid pathway, and our result suggests that it might have a role in this pathway. It is a protein of the 2-oxoglutarate (2OG) and Fe(II)-dependent oxygenase superfamily. We also find that all the genes in module 43 are HVGs. Most edges of the module are observed during the day, and the genes in the module have very similar expression patterns throughout the time course with a peak of expression at the beginning of the day and another one at the end of the day. However, none of the genes in the module 43 are also present in the averaged time course network.

Overall, we find that several modules in the variability network are functionally relevant, with modules showing enrichment for functions such as photosynthesis, DNA packaging and glucosinolate or flavonoid metabolism, even in the absence of genetic and environmental perturbations. Moreover, we could identify a potential role in the enriched pathways for some genes, based on their co-expression with other genes in the same module.

### Identification of new targets for GI, PIF4 and PRR5

To go further in the functional analysis of the modules, we looked for enrichment of targets of TFs in the modules. We focussed on TFs for which ChIP-seq were performed in similar conditions (seedlings grown in day/night cycles), and for which a list of target genes have been previously published [39–42]. This way, we identified an enrichment in modules for targets of SPL7, GI, PIF4 and PRR5 (Table S4). For example, all 41 genes in the module 8 are SPL7 targets (Table S4) [39]. This is significantly more compared to the entire network in which 244 genes are SPL7 targets (14%). SPL7 targets have been previously shown to be enriched in multiple GO terms, including photosynthesis [39], in agreement with the predominant role in photosynthesis of genes in this module 8.

We also find that 6 out of 7 genes in the module 64 are targets of GI (Table S4, Fig 4a). This is more compared to the entire network in which 394 genes are GI targets (22%). To explore more in detail GI binding at the genes in the module 64, we downloaded the ChIP-seq data, mapped it on the *Arabidopsis thaliana* genome and looked at the ChIP-seq signal for GI at all the 7 genes of the module 64 [40]. We find a strong signal for the GI ChIP-seq at the promoter of the 6 genes that were already identified as GI targets (Fig 4a). Interestingly, the signal for GI at the 7th gene, AT1G03630, not previously described as a GI target, is equally strong at the promoter of the gene (Fig 4a). This result indicates that AT1G03630 is also a target of GI, even if it has not been previously identified as such. AT1G03630, or PORC, encodes for a protein with protochlorophyllide oxidoreductase activity that is NADPH- and light-dependent [43].

**Fig 4.**
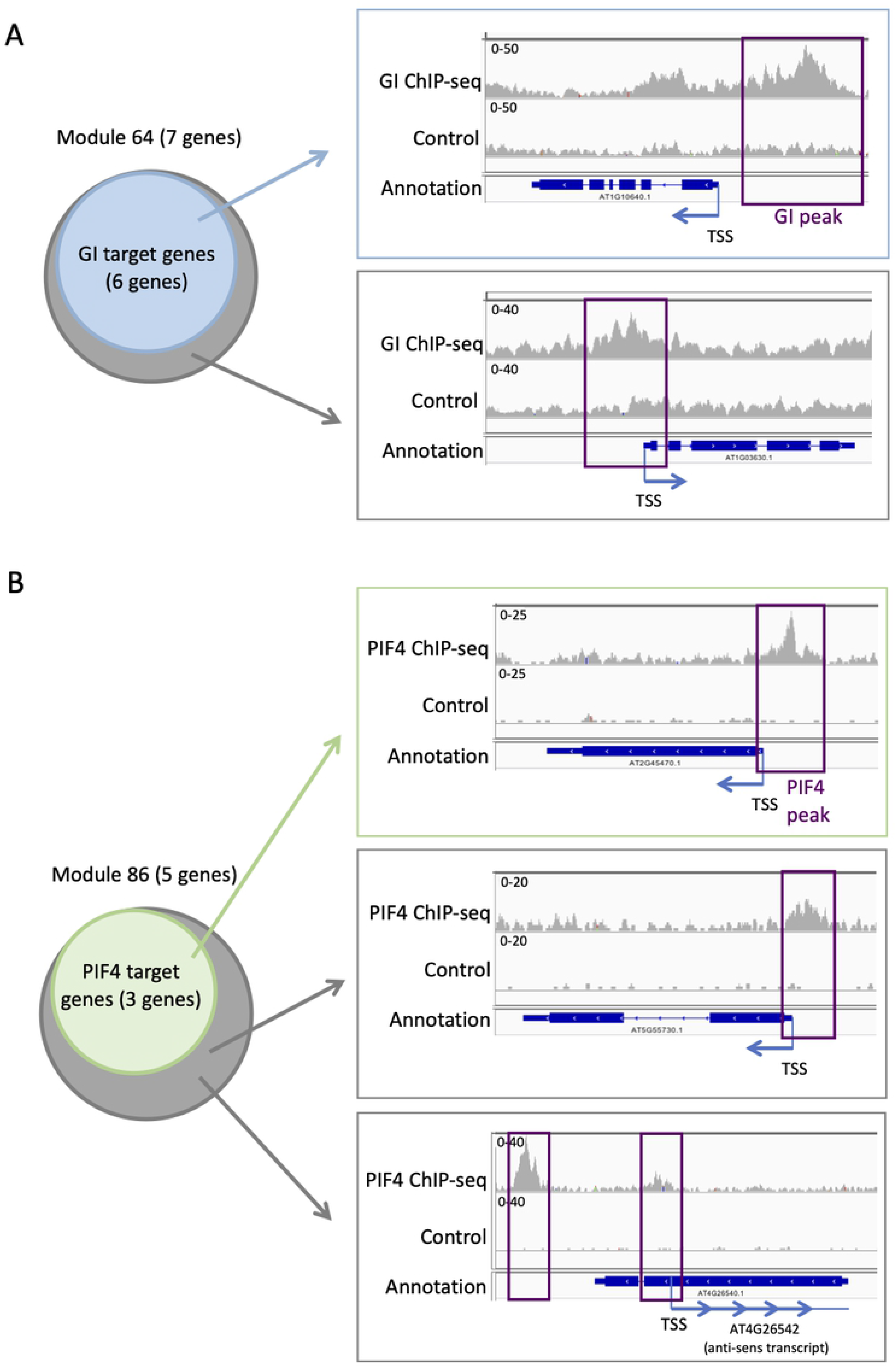
Additional TF targets can be identified using TF targets enrichment in modules. (A) Analysis of GI TF targets on the module 64: 6 of the 7 genes in the module 64 are known targets of GI (left). IGV screenshot showing the signal for the GI ChIP-seq (right) at a known GI target (top) and for the seventh gene in the module 64 that is not known as a GI target (bottom). (B) Analysis of PIF4 TF targets on the module 86: 3 of the 5 genes in the module 86 are known targets of PIF4 (left). IGV screenshot showing the signal for the PIF4 ChIP-seq (right) at a known PIF4 target (top) and for the two other genes in the module 86 that are not known as a PIF4 target (bottom).

Another TF with enriched targets in some modules is PIF4. We find that 3 out of the 5 genes (60%) in the module 86 are PIF4 targets (Table S3, Fig 4b), while only 305 genes in the full network are PIF4 targets (17.6%). To explore more in detail PIF4 binding at the genes in the module 86, we downloaded the ChIP-seq data, mapped it on the *Arabidopsis thaliana* genome and looked at the ChIP-seq signal for GI at all the 5 genes of the module 86 [41]. We observe a strong signal for the PIF4 ChIP at the promoters of the 3 known targets in the module 86 (Fig 4b). We also see a clear signal for the PIF4 ChIP for the two other genes in the module 86, AT4G26542 and AT5G55730, suggesting that they are also targets of PIF4 (Fig 4b). AT4G26542 is an anti-sens transcript for AT4G26540. AT5G55730 (FLA1) encodes a fasciclin-like arabinogalactan-protein 1 [44].

Finally, we find an enrichment for PRR5 targets in modules 8 and 21 with respectively 78% and 70% of genes in the module that are PRR5 targets [42]. For comparison, 27% of all genes in the network are PRR5 targets. To explore more in detail PRR5 binding at the genes in the module 8, we downloaded the ChIP-seq data, mapped it on the *Arabidopsis thaliana* genome and looked at the ChIP-seq signal for PRR5 at all the 41 genes of the module 8 [45]. We find a strong ChIP-seq PRR5 signal at the 32 target genes in the module 8, and a similarly strong signal for most of the other 9 genes in the module that were not listed as a PRR5 target (Fig S7a). In order to look for PRR5 targets, and to expand the analysis to other modules, we re-identified peaks for the PRR5 ChIP-seq and looked for PRR5 targets in the modules with a high proportion of already described PRR5 targets (Table S4). This way, we identified 5 additional PRR5 targets in the module 8, and 9 additional PRR5 targets in the module 21. When combining the PRR5 targets from both analyses, the total percentage of PRR5 targets is 90% in the module 8, and 83% in the module 21 (Fig S7b). These results suggest that most, if not all, genes in the module 8 and 21 are in fact PRR5 targets.

Overall, we find that some modules are enriched for TFs targets, and that this enrichment can be used to identify additional targets for the TF in the modules showing enrichment for its targets.

## Discussion

In this work, we have analysed gene co-expression networks inferred using expression data generated in the absence of genetic and environmental perturbations. To do this, we made use of an already published dataset of transcriptomes performed on single seedlings that were grown in the same environment. We showed that genes do not need a high level of gene expression variability between seedlings to be able to integrate them in the network (Table 1). Moreover we find that modules identified in this network are biologically relevant, as several are strongly enriched in GOs (Fig 3) and in TF targets (Fig 4). Based on these enrichments, we speculated that AT4G22870 could also have a role in flavonoid metabolism and identified new targets for the TFs GI, PIF4 and PRR5.

We find that it is possible to infer gene co-expression networks using transcriptomes of genetically identical plants grown in the exact same environment. This is in agreement with previous work, where co-expression networks have been inferred on transcriptomes generated on individual plants and for which genetic and environmental effects have been removed *in silico* [26]. We also find an interesting topology of the network with some modules more connected with one another, and that connected modules share similar characteristics in terms of percentage of edges detected during the day or night, and percentage of HVG (Fig 5 and S8). We observe that modules have either a high or low percentage of HVG, but rarely a mix of HVG with non HVG. This suggests that some pathways are more variable than others. We find that, in general, modules with genes involved in the response to the environment are also composed of a high percentage of HVG. This is the case for example for the module 37, enriched in photosynthesis (100% of HVG), the module 43, enriched in flavonoid metabolism (100% of HVG), or the module 1, enriched in glucosinolate metabolism (78% of HVG). Flavonoids are secondary metabolites and have been shown to be involved in many biotic and abiotic responses in plants [46]. And glucosinolates are involved in the response to pathogens [47]. In agreement with our observation, previous work showed that HVG are usually involved in the response to the environment [28,48–51]. In particular, plant-to-plant variability has already been observed for glucosinolates [30], showing that the variability in expression we observe for genes involved in this pathway can lead to differences in glucosinolate content between plants.

**Fig 5.**
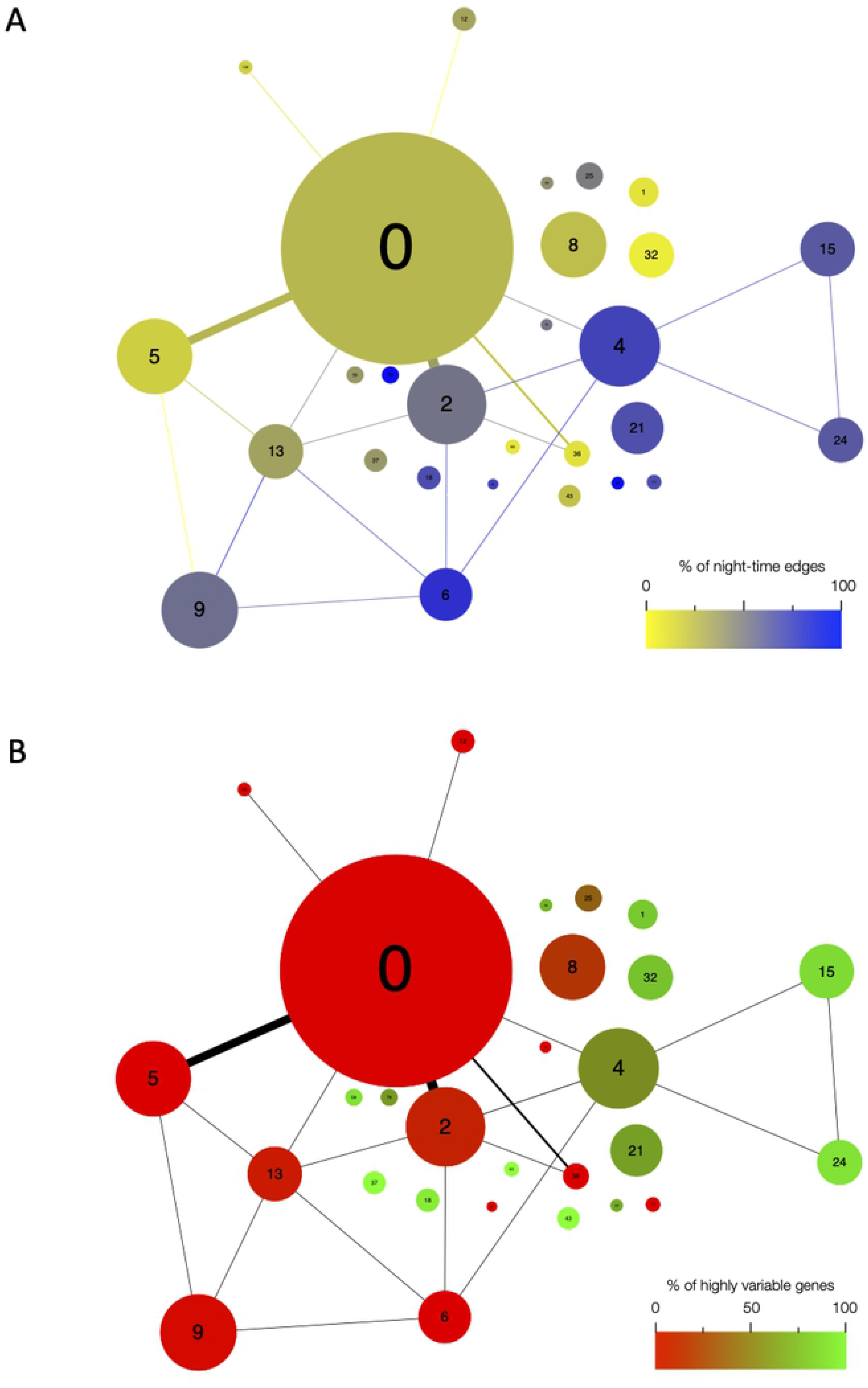
Network architecture is mainly influenced by the time of day when edges are detected, and by the presence of highly variable genes. Organisation of modules in the network, with the size of circles representing the module size (i.e. number of edges). Number of edges connecting the modules are represented by the thickness of the lines between modules. The number in each module corresponds to the module number. (A) Modules are color coded based on the percentage of edges that are detected during the night in each module. Blue modules are composed of a majority of night-time edges, while yellow modules are mainly composed of day-time edges. (B) Modules are color coded based on the percentage of highly variable genes (HVG) in the modules. Green modules are composed of a majority of HVG while red modules have a low percentage of HVG.

Like for Bhosale and colleagues, we find that the modules of the network identified in absence of genetic and environmental perturbation are biologically relevant and can be used to speculate new gene function or regulation. We only explored the function for the most obvious GO enrichment in modules as GO can be sparse for some functions and many genes do not have a GO. For example the module 43 is enriched in genes involved in the flavonoid pathway. We speculate that AT4G22870, a member of this module, is also involved in the flavonoid pathway. To support our suggestion, AT4G22870 codes for a protein of the 2-oxoglutarate (2OG) and Fe(II)-dependent oxygenase superfamily and three 2-oxoglutarate- and ferrous iron-dependent oxygenases have been previously shown to be involved in flavonoid biosynthesis [38]. Most importantly, this new potential candidate gene could not have been detected by analysing the network inferred using day/night environmental fluctuations as none of the genes in the module 43 are also present in the averaged time course network. We find several modules with enrichment for genes involved in photosynthesis. In particular, the modules 8 and 37. The main distinction between these two modules is that module 8 is composed of genes from the nuclear genome, while module 37 is mainly composed of genes from the chloroplast genome. Our approach was not designed to specifically identify and separate genes from different organelles, suggesting that genes from the nuclear and chloroplast genomes involved in photosynthesis vary differently in expression between seedlings. Our result is in agreement with the fact that organelle functional modules can be detected in *Arabidopsis thaliana* [52]. However, genes that are not from the nuclear genome are usually ignored in network analysis, and it would be of interest to integrate them in the future.

Finally, we identified enrichment for targets of the TFs GI, PIF4 and PRR5 in different modules, and used this enrichment to highlight new targets. We find that in most cases, when a module is enriched in targets for a TF, the remaining genes of that module are also targets of this TF. By reanalysing the ChIP-seq data for PRR5, we could increase the percentage of targets in modules already showing a strong enrichment. This result shows the double interest of combining co-expression networks with ChIP-seq data [21,22]. On the one hand, ChIP-seq data adds information about the regulation of genes in the co-expression network. On the other hand, the co-expression network is a good way to focus on some of the targets of the TF to better understand their regulation and also to detect extra targets. In the case of PRR5, we find that 90% of the genes in the module 8 are targets of this TF. Genes in the module 8 are involved in photosynthesis. This is in agreement with the fact that the circadian clock, of which PRR5 is a core member, has been shown to regulate the photosynthesis [53–55].

The functional characterisation of the network has been restricted to some modules with obvious GO enrichments, and to TFs for which ChIP-seq data and lists of targets were available and performed in similar conditions. However this network, being the first to be performed in absence of genetic and environmental fluctuation, could bring further information on other pathways we have not explored in this paper. Moreover, our approach could reveal co-regulations that might not be detected using environmental perturbations, as shown by the fact that the variability network provided additional co-expression relationships that were not detected in a network inferred on the same dataset using expression fluctuations caused by the day/night cycle. That is why we encourage readers to look at the modules for their genes or pathway of interest, and have developed an interactive website where readers can do so (https://jlgroup.shinyapps.io/VariabilityNetwork/).

We show that most genes in the network are not HVG (Table 1), showing that high gene expression variability between seedlings is not needed to be able to detect co-expression. These results indicate that we are not in the presence of random fluctuation in expression, or noise, but that pathways are slightly differently regulated in individual seedlings even if the plants are in the same environment. Our approach uses these small differences between seedlings that might be caused by microenvironmental fluctuation, or a different state of the plant caused by internal factors. It indicates that plants are very sensitive to minor changes in their environment, and that we could harness this sensitivity to better understand gene expression regulation. Phenotypic differences have been observed between genetically identical plants grown in the same environment [27,29,30,56–58], indicating that the changes in expression of pathways we highlight here might be physiologically relevant. It shows that it is not necessary to perform experiments in very different environmental conditions to identify co-expression networks that could be relevant to the studied pathway. Strong fluctuations (mutants, over-expressors, environmental fluctuations) could potentially affect a big part of the transcriptome that could mask some co-expressions of interest showing the usefulness of our approach in some contexts. Our work shows the interest in harnessing gene expression variability between genetically identical individual plants in order to better understand gene regulation in a context where differences between plants are not known and probably very subtle.

## Materials and Methods

### Transcriptome data

The transcriptomes we used were already published (GSE115583; [28]), and performed on single seedlings, for a total of fourteen seedlings per time point every two hours over a 24 hours cycle. Expression levels and corrected variability levels for all genes were downloaded from https://jlgroup.shinyapps.io/AraNoisy/, as this data had been already corrected as previously described [28].

### Network construction

#### Variability network

For each of the 12 time points (0h, 2h, 4h, … 22h) we calculated the Spearman correlation between every pair of genes, using their expression profiles across the 14 seedlings (Fig 1a). Using a Benjamini-Hochberg correction with a false-discovery rate of 10% the most significant correlations were selected, and further filtered by only considering those for which a significant correlation appeared in four consecutive time points (with one gap allowed, e.g. 8h, 10h, 14h, 16h). These correlations form the edges of the variability network. We also calculated a version of the network using a filter that only required three consecutive time points, and calculated network modules using the same community detection algorithm. As can be seen in Fig S1, similar modules with a similar overall connectivity between them are found, which confirms the robustness of the modules in our original network. All network analysis was carried out using the Python NetworkX and python-louvain libraries.

#### Averaged time course network

For the averaged time course network we calculated the mean expression across all seedlings for every time point, generating a time series of average expression for every gene. We again calculated the Spearman correlations for every pair of genes and generated a network by applying the Bonferroni correction as a (highly conservative) significance cutoff. This yielded a network that was similar in size to the variability network. All network analysis was carried out using the Python NetworkX and python-louvain libraries.

### Community detection

The Louvain algorithm [33] community detection algorithm was used to identify modules in the networks. This algorithm attempts to maximise the modularity of the network by searching the space of network partitions. Due to the size of the search space it is unable to find the global maximum. The composition of modules may therefore (as with most community detection algorithms) vary somewhat between runs of the algorithm.

### RT-qPCR

Col-0 WT *Arabidopsis thaliana* seeds were sterilised, stratified for 3 days at 4°C in dark and transferred for germination on solid 1/2X Murashige and Skoog (MS) media at 22°C in long days for 24 hours. Using a binocular microscope, seeds that were at the same stage of germination were transferred into a new plate containing solid 1/2X MS media. Seedlings were grown at 22°C, 65% humidity, with 12 hours of light (170 μmoles) and 12 hours of dark in a conviron reach-in cabinet. After 7 days of growth, seedlings were harvested individually into a 96-well plate and flash-frozen in dry ice. Sixteen seven-day old Col-0 WT *Arabidopsis thaliana* seedlings were harvested individually and flash-frozen in dry ice at ZT6 and at ZT14. All seedlings harvested in a given time point were grown in the same plate. Total RNA was isolated from 1 ground seedling. RNA concentration was assessed using Qubit RNA HS assay kit. cDNA synthesis was performed on 700ng of DNAse treated RNA using the Transcriptor First Strand cDNA Synthesis Kit. For RT-qPCR analysis, 0.4 μl of cDNA was used as template in a 10 μl reaction performed in the LightCycler 480 instrument using LC480 SYBR green I master. Gene expression relative to two control genes (SandF and PP2A) was measured (See Table S5 for the list of primers used for RT-qPCR).

### Gene Ontology term enrichment

We used the Ontologizer [59] command line tool with Bonferroni multiple-hypothesis correction to perform Gene Ontology (GO) term enrichment analysis of the network modules. Only the significantly enriched non redundant GO are shown.

### ChIP-seq data and analysis

ChIP-seq data were downloaded from GSE45213 for SPL7 [39], from GSE129865 for GI [40], from GSE43286 for PIF4 [41] and from GSE36361 for PRR5 [42].

ChIP-seq data were analysed in house, using a combination of publicly available software and in-house scripts. Reads were aligned to the TAIR10 genome using Bowtie2 [60]. Potential optical duplicates were removed using Picard tools (https://github.com/broadinstitute/picard). Peak calling was performed using MASC2 [61], with the corresponding INPUT used as a reference. Snapshots of ChIP-seq signal around targets were shown using the Integrative Genomics Viewer (IGV [62]).

## Authors’ contribution

SC conceived the project. SC, JCWL and SA designed the project. SA inferred the networks. SC and SA performed downstream analyses of the network and SC interpreted the data. MB performed the RT-qPCR validation. SC, JCWL and SA wrote the article, with SC writing the first draft.

## Supplementary Figures and Tables legends

**Supplementary figure 1** Expression in seedlings of genes in module 1, from the RNA-seq data, with one line per gene. Expression is mean normalised for each gene.

**Supplementary figure 2** Comparison of edges in modules detected in the networks containing edges present in 3 or 4 consecutive time points. Modules detected in the network based on edges present in at least 3 consecutive time points are shown in blue. Modules detected in the network based on edges present in at least 3 consecutive time points are shown in red. For the later, the percentage of edges of the modules that are also detected in the blue modules are indicated.

**Supplementary figure 3** Number of edges in the final network that are detected in each time point, for every module containing at least 5 genes.

**Supplementary figure 4** a. Correlation in expression between seedlings for genes of the module 1 and module 21, for the RNA-seq experiment. AT5G07690 at ZT14 was removed as it is not expressed.

b. Correlation in expression between seedlings for genes of the module 1 and module 21, based on a RT-qPCR replicate of the RNA-seq experiment. Sixteen seedlings where harvested at ZT6 and at ZT14.

c. Normalised expression level in the fourteen seedlings for the genes of the module 21, from the RNA-seq data. Expression level for AT4G13250 is shown as the x axis while expression for the other genes of the module are shown on the y axis. Expression is mean normalised for each gene.

**Supplementary figure 5** a. Inter-individual gene expression variability profiles throughout the time course for genes in each module with 5 genes or more. Each line represents the corrected variability level for one gene:: corrected CV^2^=[log_2_(CV^2^/trend)], with CV^2^=variance/(average^2^)] (see Cortijo et al., 2019).

Modules are ordered by the percentage of HVG (high to low). Modules highlighted in blue contain 75% or more of HVGs. Modules highlighted in red contain 10% or less of HVGs.

b. Heatmap of normalized gene expression for genes in modules 2, 6 (less than 15% HVG), module 4 (55% of HVG) and modules 15 and 24 (more than 85% of HVG). Expression is shown in single seedlings from the time point ZT20. Expression is mean normalised: expression in a seedling/ averaged expression in all seedlings. The left color coded bar indicates the module of each gene.

**Supplementary figure 6** a. Expression profiles throughout the time course for genes in module 8. Each line represents the normalised expression (z-score) for one gene. Genes of the photosystem I, II or the light harvesting system are in blue.

b. All edges and nodes of module 8. Genes of the photosystem I, II or the light harvesting system are in blue.

**Supplementary figure 7** a. Analysis of PRR5 TF targets on the module 8. 32 of the 41 genes in the module 8 are known targets of PRR5 (left). IGV screenshot showing the signal for the PRR5 ChIP-seq (right) at a known PRR5 target (top) and for a gene in the module 64 that is not known as a PRR5 target (bottom).

b. Comparison of published (blue) and realised (grey) PRR5 targets in modules 8 (left) and 21 (right).

**Supplementary figure 8** Organisation of modules in the network, with the size of circles representing the module size (i.e. number of edges). Number of edges connecting the modules are represented by the thickness of the lines between modules.

a. Most enriched GOs are written in each module.

b. Modules are color coded based on the percentage of edges in the averaged time course network. Dark blue modules have a high percentage of genes in the averaged time course network while light blue modules have a low percentage of genes in the averaged time course network.

**Table S1** List of genes in each module

**Table S2** GO enriched (corrected p-value<0.5) in each module

**Table S3** Function of genes in modules 1, 43 and 71

**Table S4** Number and percentage in the modules of targets for the TFs SPL7, GI, PIF4 and PRR5

